# CryptKeeper: a negative design tool for reducing unintentional gene expression in bacteria

**DOI:** 10.1101/2024.09.05.611466

**Authors:** Cameron T. Roots, Jeffrey E. Barrick

**Affiliations:** Department of Molecular Biosciences, Center for Systems and Synthetic Biology, The University of Texas at Austin, Austin, Texas 78712, U.S.A.

**Keywords:** plasmid instability, reliability and reproducibility, computational DNA sequence design, design-build-test cycle, recombinant protein overexpression

## Abstract

Foundational techniques in molecular biology—such as cloning genes, tagging biomolecules for purification or identification, and overexpressing recombinant proteins—rely on introducing non-native or synthetic DNA sequences into organisms. These sequences may be recognized by the transcription and translation machinery in their new context in unintended ways. The cryptic gene expression that sometimes results has been shown to produce genetic instability and mask experimental signals. Computational tools have been developed to predict individual types of gene expression elements, but it can be difficult for researchers to contextualize their collective output. Here, we introduce CryptKeeper, a software pipeline that visualizes predictions of bacterial gene expression signals and estimates the translational burden possible from a DNA sequence. We investigate several published examples where cryptic gene expression in *E. coli* interfered with experiments. CryptKeeper accurately postdicts unwanted gene expression from both eukaryotic virus infectious clones and individual proteins that led to genetic instability. It also identifies off-target gene expression elements that resulted in truncations that confounded protein purification. Incorporating negative design using CryptKeeper into reverse genetics and synthetic biology workflows can help to mitigate cloning challenges and avoid unexplained failures and complications that arise from unintentional gene expression.

## Introduction

RNAs and proteins may be unexpectedly transcribed and translated from a DNA sequence. This type of cryptic gene expression can complicate studying and engineering biological systems. Cryptic gene expression can occur when natural DNA sequences contain promoters and ribosome-binding sites that are not annotated because they are redundant, antisense, or internal to genes. It can also emerge when sequences are moved into a new cellular context (e.g., cloning eukaryotic sequences in *E. coli*) (1–4) or as a consequence of engineered changes to sequences (e.g., combining genetic parts, optimizing codon usage, or introducing artificial watermarks) (5,6). Cryptic gene expression products that interfere with the intended function of an engineered DNA construct may cause a design to be deemed a failure.

Worse yet, cryptic gene expression may occur unbeknownst to researchers, causing them to misinterpret experimental results (7,8). Unintentional expression of genes, truncated pieces of genes, or out-of-frame products is often burdensome or even toxic to a host organism, creating a strong selection pressure favoring cells with mutations in the engineered DNA sequence (1–4,9). Rapid evolution of escape mutants that eliminate these or other sources of burden can be one reason that certain sequences are unreliable or even unconstructable (10,11).

Both the process and products of gene expression can be burdensome to a cell. Studies of recombinant protein overexpression have shown that growth rates of bacterial cells decrease in proportion to how much of their translational capacity, usually determined by the number of ribosomes, is redirected to expressing exogenous proteins (12,13). Expression of some proteins is also directly deleterious due to their activities, whether they are enzymes that rewire metabolism in ways that redirect limiting resources away from cellular replication or disrupt cellular homeostasis in other ways (11,12). It is rarer for transcription of RNA alone to cause an appreciable burden on a bacterial cell, but it has been documented in yeast protein overexpression systems (14,15).

Negative design is the process of eliminating undesirable qualities to engineer a safer or more effective system (16). The relative rates at which ribosomes initiate translation from different start codons in *E. coli* and other bacteria can be accurately predicted from characteristics of their ribosome-binding sites and surrounding sequences (17–19). Therefore, one negative design strategy for solving problems stemming from cryptic protein expression is to concentrate on redesigning a DNA sequence to eliminate the potential for unwanted translation, whether or not there is any evidence that a relevant mRNA is transcribed. Because translation and transcription are coupled in bacteria, disrupting translation is also expected to reduce RNA levels by short-circuiting transcription and promoting mRNA degradation (20,21).

Another negative design approach would be to eliminate cryptic transcription so mRNAs are not produced in the first place. Unfortunately, tools for predicting bacterial promoters and terminators currently have limited accuracy, only make qualitative predictions, and/or are not openly accessible (22–40). Furthermore, many promoter prediction tools use coding sequences as their non-promoter group during development, an assumption which could lead to systematically misclassifying cryptic promoters found within ORFs (30,37,40). Even so, predictions of promoters and terminators could provide additional context for interpreting predictions of translated reading frames and warn of other potential problems with a sequence design, such as the accidental production of inhibitory transcripts that are antisense to known genes.

Here we describe CryptKeeper, an open-source software tool that integrates and displays predictions of *Escherichia coli* gene expression elements in engineered DNA constructs, such as plasmids. CryptKeeper is designed to allow users to evaluate the potential for cryptic gene expression that may interfere with the construction or function of a DNA sequence. We demonstrate the utility of CryptKeeper by using it to analyze the results of several prior studies in which researchers identified cryptic gene expression that was problematic and then redesigned their sequences to avoid it.

## Methods

### Software overview

CryptKeeper integrates the output of several tools that predict bacterial gene expression elements from DNA sequences (**Fig. 1A**). It accepts input sequences in GenBank or FASTA format. It displays predictions of translation initiation sites, promoters, Rho-dependent terminators, and intrinsic (Rho-independent) terminators. These predictions are summarized with a translational burden score and displayed in an interactive visualization so that a user can evaluate whether there is the potential for cryptic gene expression that may interfere with the function or stability of their DNA sequence. CryptKeeper is a Python package. It and all of its dependencies can be installed as Bioconda packages.

**Fig. 1.**
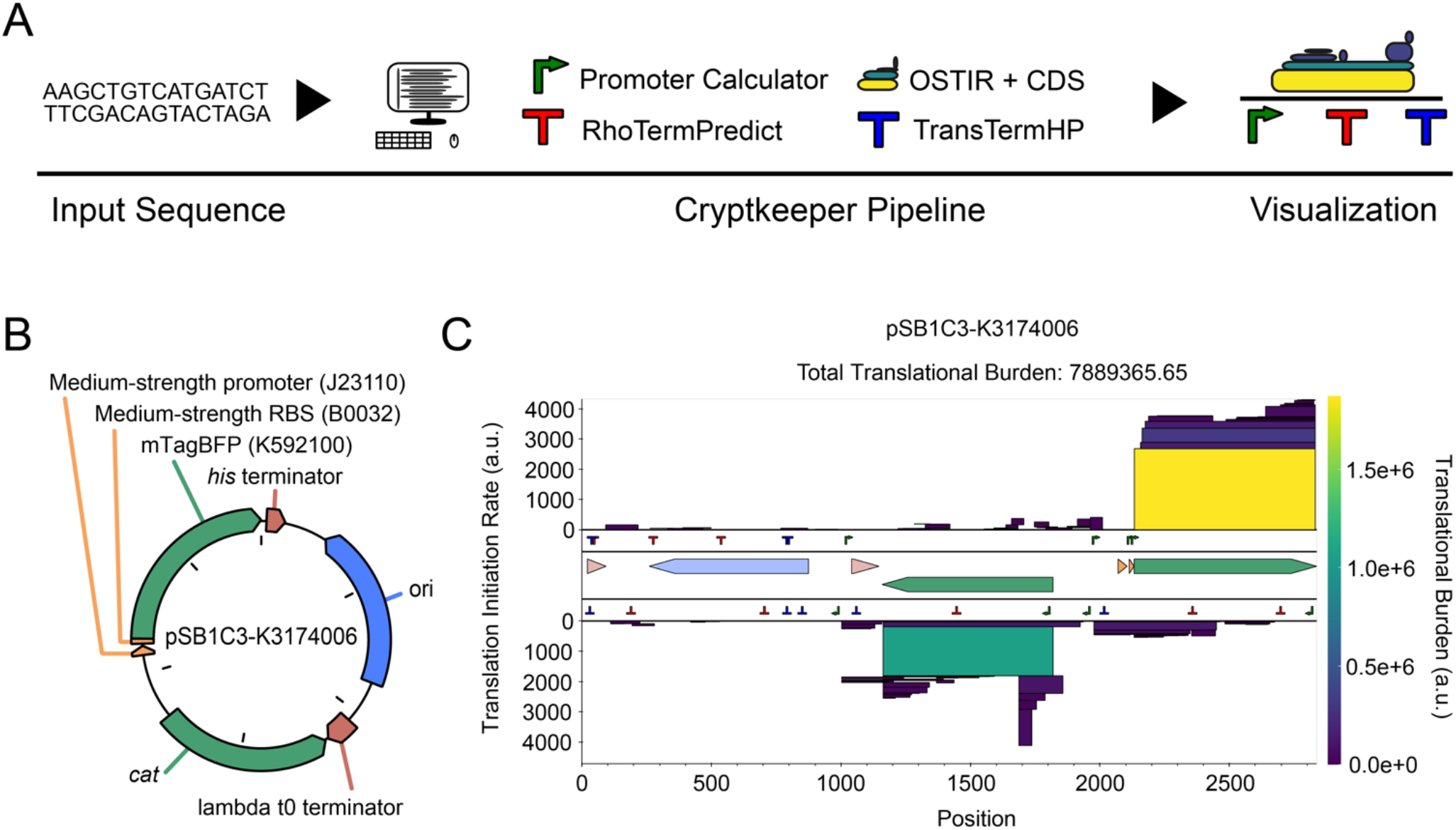
(A) CryptKeeper overview. Predictions of gene expression signals in an input DNA sequence are integrated into an interactive plot and used to calculate an overall translational burden score. (B) pSB1C3-K3174006, an example of a plasmid engineered to express a transgene. It contains a pUC origin of replication, chloramphenicol resistance cassette, and BioBrick K3174006, which expresses mTagBFP (K592100) under control of a medium-strength constitutive promoter (J23110) and a medium-strength ribosome binding site (B0032) (10). Figure adapted from pLannotate output (41). (C) CryptKeeper output for pSB1C3-K3174006. The outermost tracks display protein-coding sequences on the forward and reverse strands as stacked boxes with heights proportional to their predicted translation initiation rates and colors and areas proportional to their individual translational burden scores. The next inner two tracks display predictions of RNA expression signals on each strand: promoters (green), Rho-dependent terminators (red), and intrinsic terminators (purple). The central track displays annotations from the input GenBank file (matching panel B).

### Translational Burden Prediction

Thermodynamic models can accurately predict the relative rates at which ribosomes initiate translation from different start codons (17–19). CryptKeeper calculates the translation initiation rates at all start codons in the input sequence using OSTIR version 1.1.2 (Roots et al., 2021). The translational burden of a DNA construct on an *E. coli* host cell is expected to be proportional to the number of ribosomes bound to new mRNAs transcribed from it (12,13). If rates of translation elongation and termination are fast relative to initiation and uniform, then ribosome occupancy of a given open-reading frame (ORF) will be directly proportional to the rate of initiation at its start codon and its length. Therefore, we summarize these results as a translational burden score for each ORF that is the product of its predicted translation initiation rate and its length in base pairs. Very short ORFs (<45 bases) are unlikely to contribute much to the overall burden of a construct. They are predicted by Cryptkeeper but are not shown in its graphical output (to avoid unnecessary visual clutter) when using the default settings.

### Promoter and Terminator Prediction

CryptKeeper displays predictions of *E. coli* σ^70^ promoters from a fork of the Promoter Calculator version 1.2.2 (29) that we created to add multithreading, reduce memory usage, and make it installable as a Bioconda package. Intrinsic (Rho-independent) terminators are predicted using TransTermHP version 2.09 (28). Rho-dependent terminators are predicted using a fork of RhoTermPredict version 3.4.0 (23) that we created to make it installable as a Bioconda package. CryptKeeper does not attempt to integrate transcription predictions into an overall score because these they are less accurate and complete than translation predictions. For example, the Promoter Calculator only predicts σ^70^ promoter initiation strength with a coefficient of determination of 0.45 for a test set of plasmid-encoded promoters in *E. coli* (29). While some tools exist that predict promoters that use alternative sigma factors, they are classifiers that do not quantitatively predict strength or are not open-source tools that can be run at the command line (30,37,40). By contrast, prediction of intrinsic terminators reaches >90% accuracy and specificity (28) and should generalize to many other bacterial species. However, there is almost always some read-through of these terminators (42,43) and this characteristic is not predicted by current tools. Predictions of Rho-dependent terminators may also generalize across bacteria, but they tend to have indistinct boundaries and even less is known about how well their presence and efficiencies are predicted by current algorithms (Di Salvo et al., 2019). Despite these current shortcomings, predictions of transcription initiation and termination elements may provide additional context to the user and may be sufficient for spotting problems in certain cases.

### Output

For an input DNA construct (**Fig. 1B**), CryptKeeper uses the Bokeh Python library (44) to output an HTML document that includes an interactive plot (**Fig. 1C**). This plot displays stacked boxes associated with different ORFs. The height of each box is proportional to the predicted rate of translation initiation at the start codon of the ORF, which makes its area proportional to the translational burden score. Boxes are also colored according to their burden scores on a linear scale. Promoter and terminator predictions are shown on two inner tracks, one for each DNA strand. The number of these predictions shown can be adjusted by the user. The default is to show the three strongest promoters, Rho-dependent terminators, and intrinsic terminators per kilobase of the input sequence. If a GenBank file was used as the input, features annotated in this file are shown in the central track. The CryptKeeper plot can be zoomed and rescaled, and it displays information about each predicted feature and annotation on mouseover. To facilitate further analysis by users, a table describing each predicted element is provided below the plot and in a separate comma-separated values (CSV) output file.

### Test Datasets

We tested CryptKeeper on DNA sequences from six published studies that encountered and characterized cryptic gene expression from plasmids in *E. coli* (**Table 1**). In each case, the relevant sequences were recreated *in silico*. When sufficient information was available, the entire plasmid sequences were reconstructed and analyzed. All of these studies report how researchers introduced mutations that resolved their issues, which allowed us to further examine how well CryptKeeper output tracks with the experimentally validated outcomes of redesigning DNA sequences. Plasmid annotations were based on GenBank records (45), pLannotate predictions (41), and descriptions in the relevant studies.

**Table 1.**
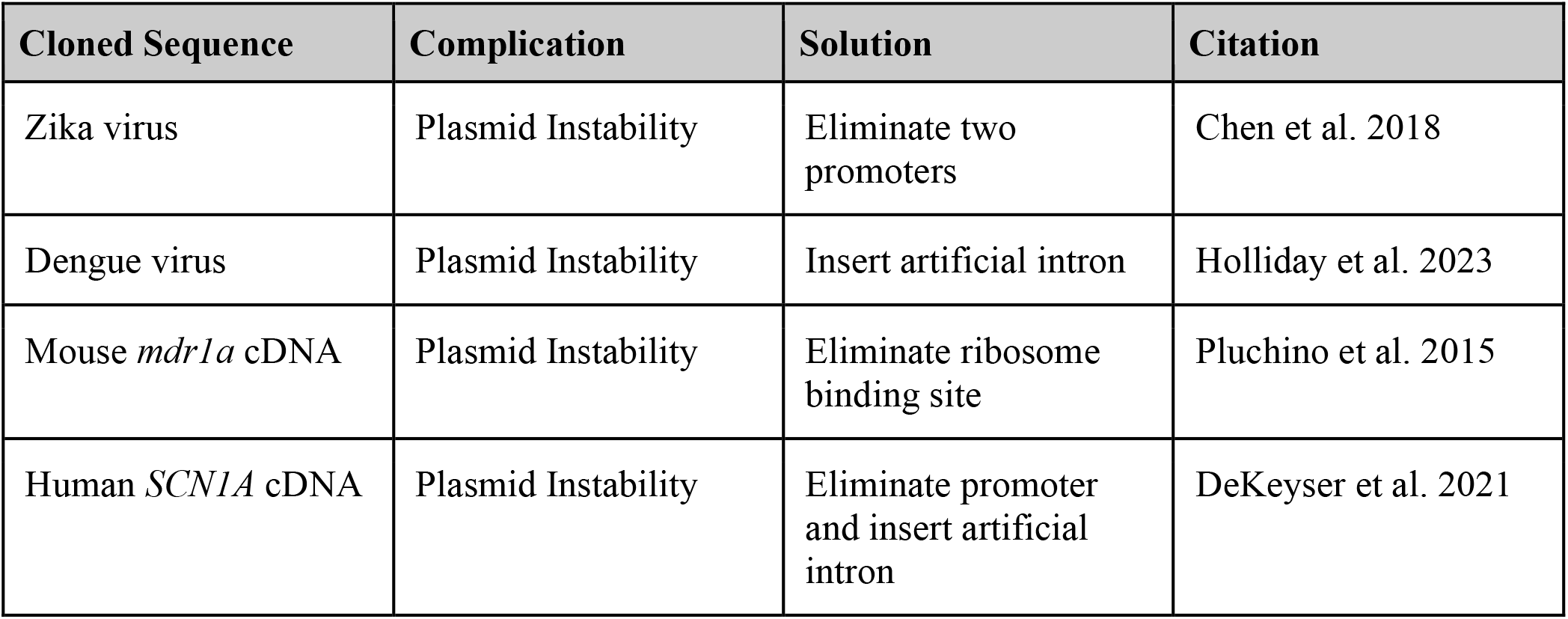
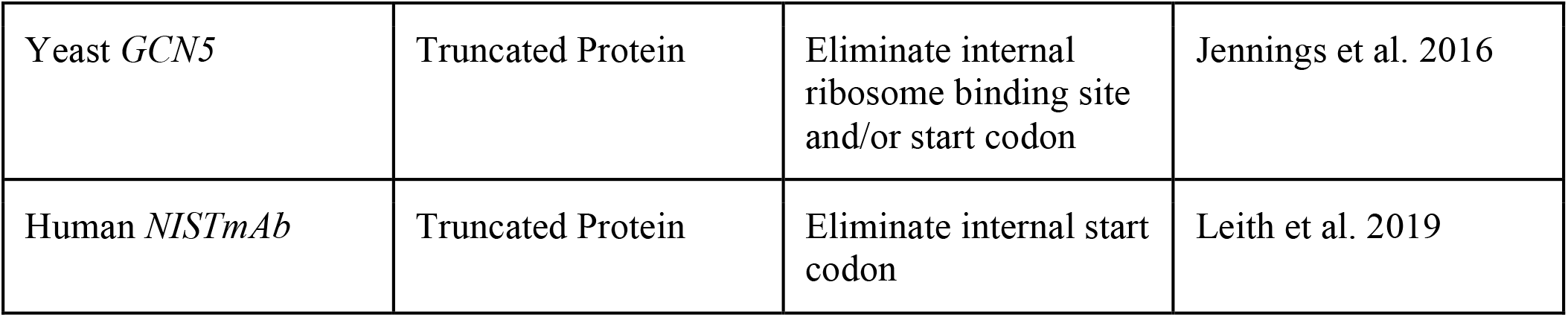
Test Datasets.

| Cloned Sequence | Complication | Solution | Citation |
| --- | --- | --- | --- |
| Zika virus | Plasmid Instability | Eliminate two promoters | Chen et al. 2018 |
| Dengue virus | Plasmid Instability | Insert artificial intron | Holliday et al. 2023 |
| Mouse *mdr1a* cDNA | Plasmid Instability | Eliminate ribosome binding site | Pluchino et al. 2015 |
| Human *SCN1A* cDNA | Plasmid Instability | Eliminate promoter and insert artificial intron | DeKeyser et al. 2021 |
| Yeast *GCN5* | Truncated Protein | Eliminate internal ribosome binding site and/or start codon | Jennings et al. 2016 |
| Human *NISTmAb* | Truncated Protein | Eliminate internal start codon | Leith et al. 2019 |

## Results

### Virus Infectious Clone Case Studies

Cloning a plant or animal virus into an *E. coli* vector makes it possible to use standard molecular biology workflows to modify its sequence. These infectious clone plasmids are a widespread reverse genetics tool for studying viruses and their applications in biotechnology. Because the virus DNA in an infectious clone is replicating in a cell that is evolutionarily distant from its eukaryotic host, it should not produce toxic virus proteins or infectious particles. However, sometimes there are cryptic gene expression elements in virus sequences that direct transcription and translation in *E. coli* cells. These products can cause significant translational burden or toxicity, leading to plasmid instability.

In a study that cloned Zika virus into a bacterial plasmid to create an infectious clone, researchers identified two putative *E. coli* promoters they designated ECP1 and ECP2 within the nucleotide sequence of the E envelope protein that they suspected were responsible for the expression of toxic products (1). CryptKeeper predicts ECP1 as the strongest promoter within the Zika genome. ECP2 is also identified by CryptKeeper, but it is among the weaker promoter predictions in the construct, so it is not shown in the plot with the default settings. The researchers found that introducing point mutations in both ECP1 and ECP2, as well as ECP1 alone, allowed them to stabilize the infectious clone. Their sequence changes reduced the transcription initiation rate predicted by CryptKeeper for ECP1 by 67% and did not change the predicted rate of ECP2 (**Fig. 2A**). Both promoters are located approximately 1800 bases upstream of an ORF with a predicted translational burden that is similar to what we found explained instability in other case studies. However, these researchers did not test for cryptic translation, so we cannot determine whether the instability they observed was due to translational burden or a toxic effect of a protein product.

**Fig. 2.**
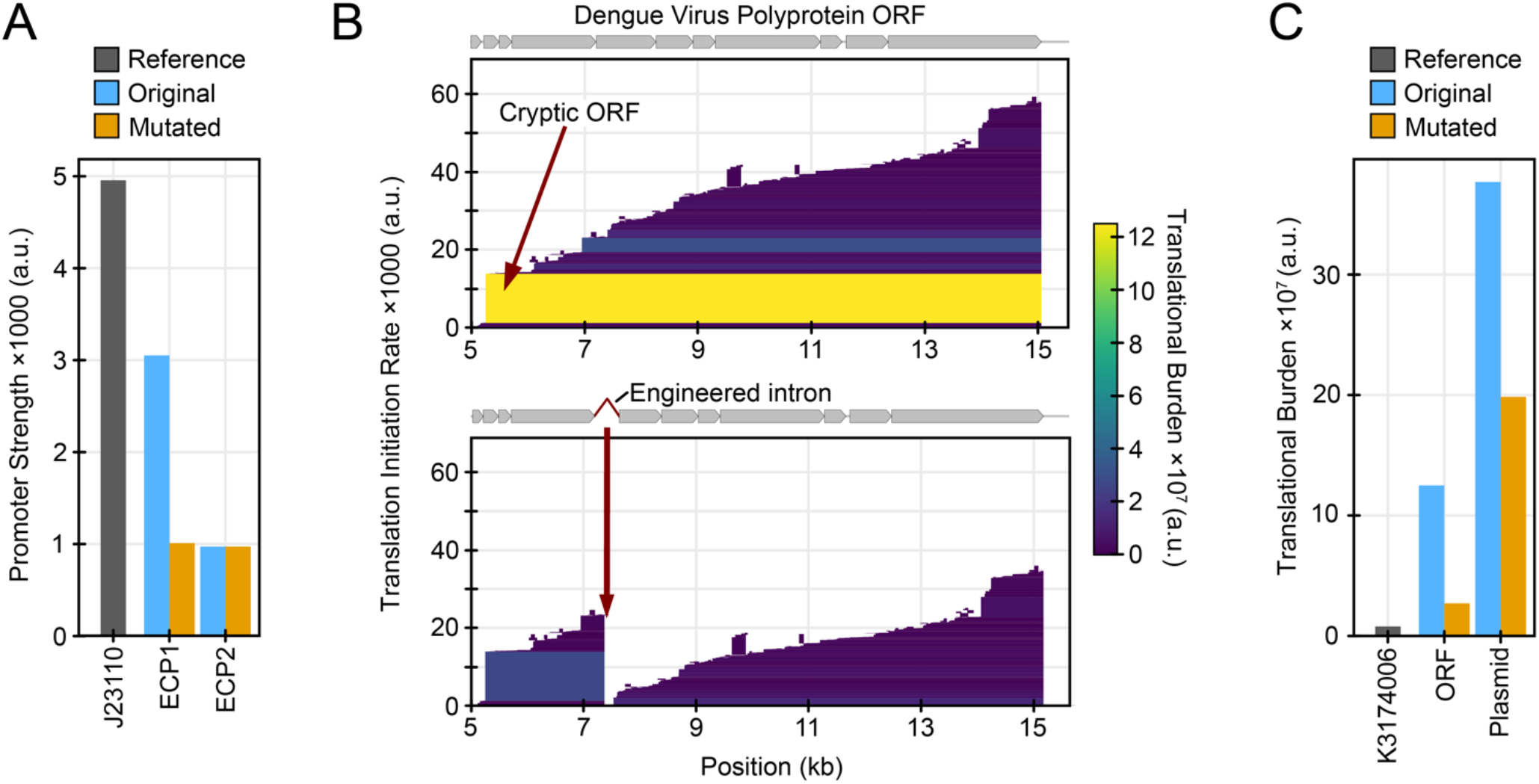
Case studies of virus infectious clone redesign. (A) Predicted strengths of promoters in a Zika virus infectious clone plasmid before and after redesign, compared to the promoter in BioBrick K3174006. (B) CryptKeeper burden plots for the Dengue virus sequence in an infectious clone plasmid before and after adding an intron that disrupts the ORF that makes the largest contribution to burden. (C) Predicted translational burden of a Dengue virus infectious clone before and after redesign compared to the predicted burden of the mTagBFP ORF from BioBrick K3174006.

In another study, researchers found that a Dengue virus infectious clone plasmid was unstable in *E. coli*, which they attributed to cryptic bacterial promoters within its 5’ untranslated region (UTR) (46). Subsequent studies produced stabilized Dengue virus infectious clones through a variety of approaches. Several copies of the TetR binding site tetO were sufficient for its binding to the 5’ UTR to prevent transcription initiation from the upstream promoters (47). Using a low-copy bacterial artificial chromosome instead of a high-copy plasmid also stabilized an infectious Dengue virus clone, presumably because this reduced all cryptic expression (47,48). Recently, researchers constructed a stable infectious clone in a high-copy plasmid by introducing a synthetic intron within the NS1 coding region (3). This addition interrupts translation of the long Dengue polyprotein in *E. coli* cells where it is not spliced, but the intron is removed and the virus RNA becomes infectious when transfected into mammalian cells. CryptKeeper predicts several promoters in the Dengue 5’ UTR, including a weak promoter overlapping with previously predicted ones. It also predicts a strong *E. coli* ribosome-binding site that initiates translation beginning at M126 of the Dengue polyprotein (**Fig 2B**). Adding the synthetic intron introduces a stop codon that interrupts this very long reading frame (**Fig 2B**), which reduces the predicted translational burden from this ORF by 88.4% and reduces the total translational burden of the complete infectious clone plasmid by 48.6% (**Fig. 2C**).

### Eukaryotic Transgene Case Studies

Gene expression costs are not restricted to plasmids that encode viruses. Individual proteins or protein complexes are often cloned into bacteria to study their functions or for biomanufacturing. Long or toxic ORFs in these constructs can present cloning issues similar to those of the infectious clone plasmids.

When cloned in *E. coli*, the mouse *mdr1a* cDNA was found to contain a bacterial promoter and ribosome binding site near the 5’ end of its ORF (4). These elements contributed to instability.

Mutating the start codon associated with the ribosome binding site (RBS) resulted in a stable plasmid. CryptKeeper predicts both the cryptic promoter and the cryptic RBS reported in the study. The researchers eliminated cryptic translation in *E. coli* by changing the M107 ATG start codon of *mdr1a* to CTG. CryptKeeper predicts that this edit should completely abolish expression of the highly burdensome ORF (**Fig. 3A, B**), thereby reducing the burden of the cDNA portion of the plasmid by 60.9% (**Fig 3D**).

**Figure 3.**
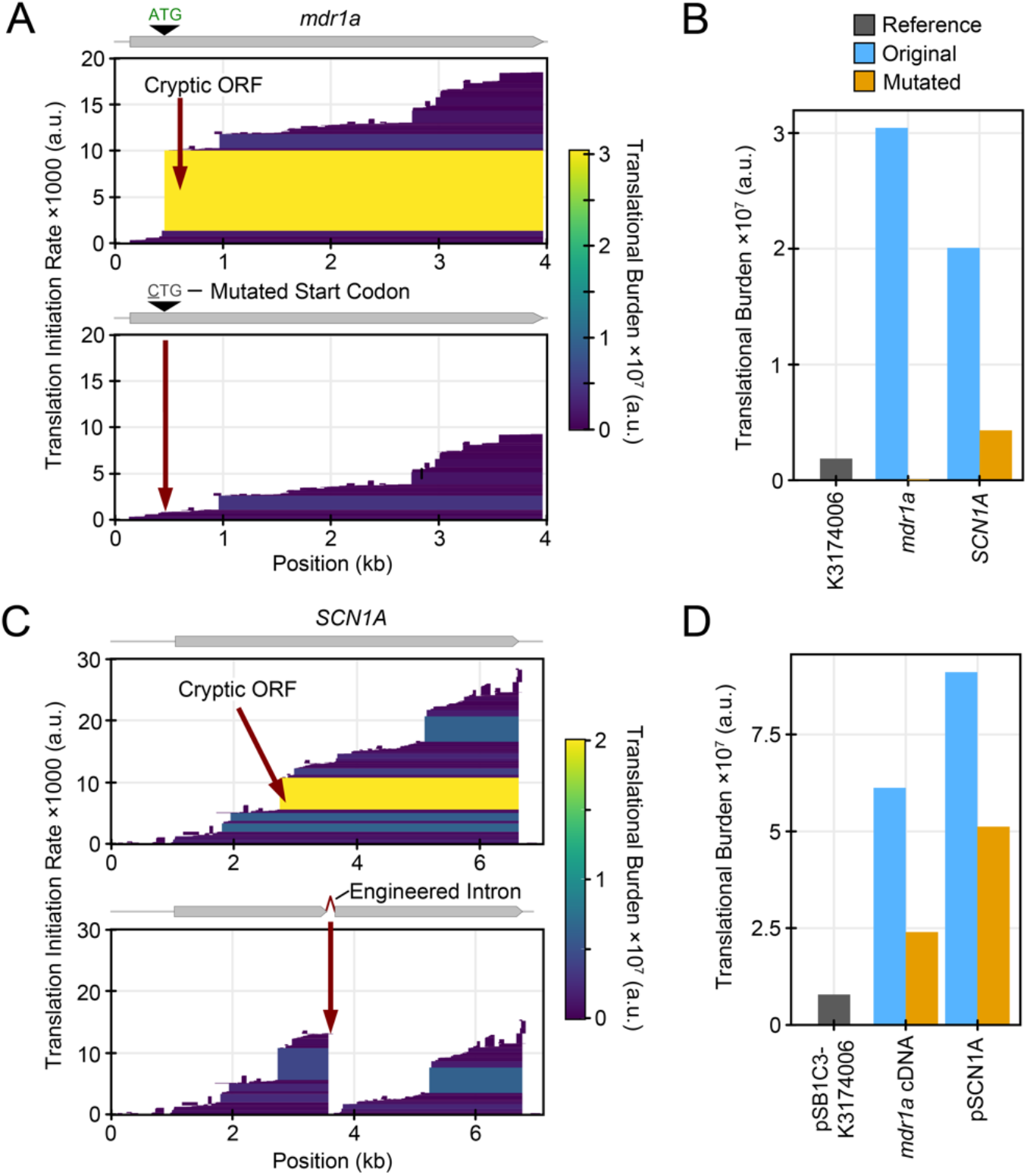
Case studies of eliminating cryptic translation from eukaryotic transgenes. (A) CryptKeeper translation predictions for a cloned mouse *mdr1a* cDNA sequence before and after mutating the start codon associated with a cryptic ORF. (B) CryptKeeper burden predictions for the mTagBFP ORF from BioBrick K3174006, the cryptic ORF of *mdr1a* before and after mutating its start codon, and the cryptic ORF of *SCN1A* before and after introducing an engineered intron. (C) CryptKeeper translation predictions for a human *SCN1A* cDNA sequence before and after redesigning it to include an engineered intron. (D) Predicted total burden of plasmid pSB1C-K3174006, the complete *mdr1a* cDNA before and after mutating the cryptic ORF start codon, and the full plasmid encoding *SCN1A* before and after introducing the engineered intron.

Another similar study found that the *SCN1A* cDNA encoding the human sodium channel Na_v_1.1 contains a cryptic promoter and translation initiation site that results in strong expression of a truncated product in *E. coli* (2). Introduction of a β-globin/IgG chimeric intron containing an in-frame stop codon was used to disrupt *E. coli* translation and establish plasmid stability in this case. CryptKeeper detects both the promoter and translation initiation site suspected of causing cryptic gene expression (**Fig. 3C**). Interruption of the cryptic ORF by the introduced intron reduces the predicted translational burden associated with its initiation site by 78.4% (**Fig. 3B**) and the score for the complete plasmid by 44.3% (**Fig. 3D**).

### Protein Truncation Case Studies

Experimental complications from cryptic translation are not limited to instability from burden. Truncated proteins produced by internal translation initiation sites can disrupt fusions to purification tags, antibody epitopes, or fluorescent reporters, decoupling these sequences from the protein of interest. It has been demonstrated that mutating predicted translation initiation sites can be used eliminate the unintentional production of truncated proteins (49). CryptKeeper is able to detect and visualize these internal ribosome binding sites, which can help researchers diagnose and redesign their sequences to prevent the unintentional translation of truncated proteins.

Researchers who cloned the yeast *GCN5* gene in *E. coli* to express and purify the yeast SAGA histone acetyltransferase observed a smaller product that was suspected to be a proteolytic degradation product of full-length SAGA on SDS-PAGE gels (7). Editing the construct to disrupt an internal RBS or start codon reduced or eliminated this band, revealing that the smaller product was a truncated protein resulting from cryptic translation. CryptKeeper predicts that the rate of translation initiation at this internal start codon is ∼350% than that of the upstream start codon for the full-length protein **(Fig. 4A)**. The mutated RBS sequence used in the study reduced the truncated protein’s predicted expression to just 8% of the unmodified truncation and 29% of the full-length protein. As shown in the study, mutating this start codon from GTG to GTT entirely eliminated the truncated product.

**Figure 4.**
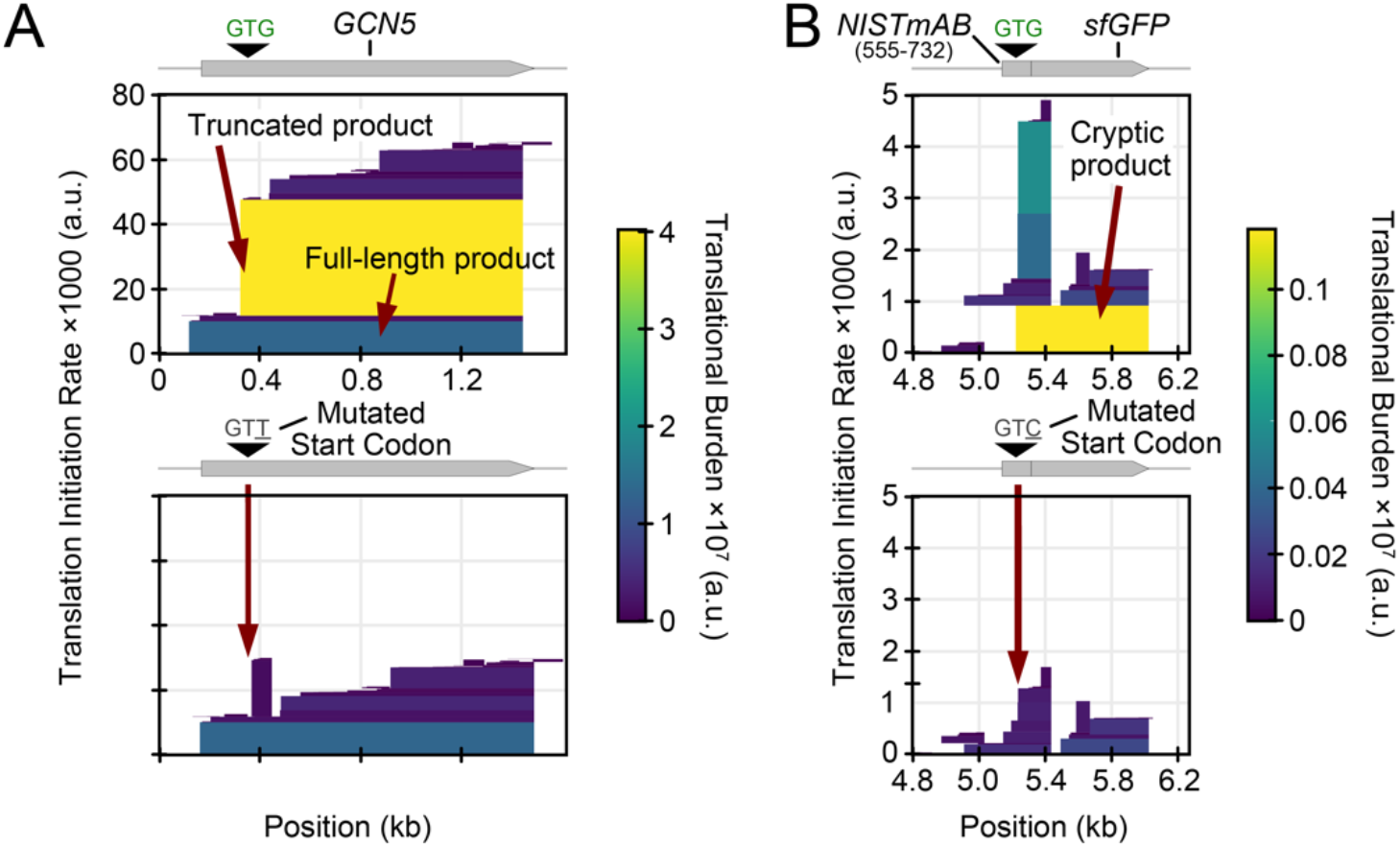
Case studies of redesign to eliminate truncated protein expression. (A) CryptKeeper translation predictions for a *GCN5* expression cassette for producing yeast SAGA histone acetyltransferase. Predictions are shown before and after mutating an internal start codon. (B) CryptKeeper translation predictions for a construct consisting of a fragment of the codon-optimized antibody NISTmAB ORF (nucleotides 555 to 732) placed upstream of a sfGFP reporter. Predictions are shown before and after mutating an internal start codon in the NISTmAB ORF.

A NISTmAb human antibody gene sequence for *E. coli* expression produced a shorter protein that copurified with the complete antibody (6). Later, it was demonstrated that this product was a truncated heavy chain produced by an RBS and GTG start codon that were unintentionally introduced during the codon optimization process (8). In a construct in which the researchers used this putative RBS and GTG codon to drive sfGFP expression to investigate the source of this unwanted product, mutating the start codon to GTC fully eliminated fluorescence. CryptKeeper identifies the unintended RBS in the sfGFP expression construct and correctly predicts that no full-length protein will be produced after the GTG to GTC start codon mutation (**Fig. 4B**).

## Discussion

DNA synthesis, assembly, and cloning workflows are critical for a wide variety of bioengineering tasks, including vector construction, protein purification, enzyme engineering, genetic circuit design, and more. Cryptic gene expression can disrupt these workflows and obfuscate experimental results, leading to abandoning constructs, time-consuming troubleshooting, or incorrect conclusions. Since many cloning failures go unexplained and unpublished, problems with cryptic gene expression are undoubtedly underreported. Currently, there is no freely accessible, open-source solution for integrating output from the ecosystem of tools for predicting gene expression elements into a visual dashboard that makes potential design issues immediately evident to a researcher. We show that CryptKeeper can effectively diagnose these issues, as described in the troubleshooting case studies.

Ultimately, the utility of CryptKeeper is limited by the computational tools that are available for predicting gene expression. For *E. coli*, these challenges currently include high rates of false-positives/false-negatives and poor quantitative accuracy when predicting promoters, even with state-of-the-art algorithms trained on large sets of experimental data (22,29,30,33,37,40). Tools for predicting transcription initiation rates driven by alternative sigma factors, transcription driven by T7 RNA polymerase, and quantitative predictions of terminator read-through are also needed to complete the picture of transcription in this host. We expect tools for predicting gene expression to improve as laboratory automation makes more extensive training sets available and as new machine learning approaches are adopted (e.g., large language models) (50,51). Ideally, it would be possible to tailor these tools for different bacterial hosts used as alternative chassis for cloning (e.g. *Vibrio natriegens*) (52) or for specific bioengineering applications (e.g., *Pseudomonas putida*) (53). Another goal for the field should be to extend these approaches to widely used eukaryotic chassis, such as budding yeast (*Saccharomyces cerevisiae*), where cryptic gene expression also poses challenges (54).

The main summary output from CryptKeeper is a translational burden score for each ORF in the input sequence. This score reflects, in relative terms, how much of a cell’s capacity for translation is expected to be redirected to this ORF, as this has been shown to be the major cause of burden for many constructs (10,12,13). The translational burden score is currently calculated simply as the translation initiation rate multiplied by the ORF length. More detailed models could account for how rare codons that slow translation or mRNA structures that act as pause sites exacerbate this burden by leading to more ribosomes than expected from the simple model becoming sequestered on certain mRNAs. This effect has been experimentally demonstrated by comparing constructs with rare codons early versus late in a reading frame (12). Incorporating these refinements into CryptKeeper’s score could be especially important for evaluating burden from cloning eukaryotic sequences with very different codon usage into *E. coli* plasmids, as is the case when constructing virus infectious clones.

CryptKeeper is most useful as a tool for negative design. In this paradigm, one takes care to avoid issues that could arise from off-target interactions when engineering a system. Other examples of negative design in synthetic biology include adding genetic insulators between modules (55), avoiding crosstalk between metabolic pathways (56), and editing DNA sequences to remove mutational hotspots (57,58). In the context of negative design, it is fine to overpredict problems, as long as alternatives without these potential issues exist in the space of possible designs. Biological sequences have so many degrees of freedom that this is often the case. For example, one can eliminate potential ribosome binding sites by making synonymous changes in one or a few codons or eliminate a start codon with single amino acid substitution. Researchers can—and we argue, should—take precautionary steps to avoid off-target translation and unintentional translational burden even if it is unclear whether any RNA containing an ORF will be transcribed. Future versions of CryptKeeper could automate the redesign of sequences for this objective. Natural gene sequences have experienced selection against off-target gene expression elements in their original biological contexts (59,60). CryptKeeper can help researchers follow suit and apply negative design to sequences they create *de novo* or transplant into new contexts to improve the reliability and reproducibility of synthetic biology.

## Material Availability Statement

CryptKeeper is open-source software released under a GPL-3.0 license. Source code, instructions, and example data are available at https://github.com/barricklab/CryptKeeper. The current version of the repository has been archived on Zenodo (DOI:10.5281/zenodo.13308762). Additionally, CryptKeeper can be installed as Bioconda package.

## Data Availability Statement

Data used for testing CryptKeeper is available at https://github.com/barricklab/CryptKeeper. The current version of the repository has been archived on Zenodo (DOI:10.5281/zenodo.13308762).

## Acknowledgements

We thank Sean Leonard and Peng Geng for helpful discussions and software testing. We also thank Davy Wybiral for helping to resolve a Python Package Index naming conflict.

## Author Contributions

Conceptualization: Cameron T. Roots, Jeffrey E. Barrick.

Funding Acquisition: Jeffrey E. Barrick.

Methodology: Cameron T. Roots, Jeffrey E. Barrick.

Software: Cameron T. Roots.

Visualization: Cameron T. Roots.

Writing – Original Draft Preparation: Cameron T. Roots, Jeffrey E. Barrick.

Writing – Review & Editing: Cameron T. Roots, Jeffrey E. Barrick.

